# Habitat use of rewilded horses and cattle as related to the functional and structural composition of plant communities in a European restored wetland ecosystem

**DOI:** 10.1101/2024.07.27.605379

**Authors:** Lilla Lovász, Fränzi Korner-Nievergelt, Valentin Amrhein

**Affiliations:** Research Station Petite Camargue Alsacienne, 1 Rue de la Pisciculture, 68300 Saint Louis, France; Department of Environmental Sciences, Universität Basel, Vesaglasse 1, 4051 Basel, Switzerland; Oikostat GmbH, Rothmättli 16, 6218 Ettiswil, Switzerland

## Abstract

1. Rewilding initiatives in European open and semi-open lowlands increasingly involve cattle and horses for ecological restoration, especially in wetland areas of high conservation value. These large herbivores contribute to spatial heterogeneity and enhance biodiversity by shaping ecosystems through movement, grazing, and resting behaviours. However, the effect of their site-specific habitat use patterns on plant communities remains unclear.
2. In this study, we investigated the fine-scale spatiotemporal distribution of rewilded cattle and horses in a recently restored alluvial grassland in a French nature reserve. We explored differences in habitat use between the two species during summer and winter on a macrohabitat scale and examined structural and functional changes in vegetation traits over four years, focusing on plant height, patch cover, species richness, and light preference, nutrient-tolerance, and mowing/grazing tolerance of plants. The study site, a former agricultural area converted into a restored alluvial nature conservation site, allowed observation of ecological processes from a “zero state”.
3. Our results suggest that cattle and horses exhibit similar habitat use with seasonal variations, potentially indicating partially shared feeding niches. The mixed-species grazing prevented vegetation overgrowth by keeping plant cover and vegetation height under control, yet without causing destructive impacts. The two herbivore species induced a clear increase in grazing-tolerant plants and slight changes in the abundance of light-preferring and nutrient-tolerant species.
4. *Synthesis*: Overall, we found that the varying spatiotemporal distribution of rewilded horses and cattle likely induces changes in plant community on the patch scale but results in vegetation stability on the landscape scale, which is known to facilitate ecosystem functioning. Our study therefore informs managers of conservation initiatives, proposing rewilding with year-round grazing horses and cattle a promising strategy for ecological restoration and natural habitat maintenance in wetland areas.

## Introduction

Rewilding with domestic cattle and horse breeds is becoming increasingly widespread in ecological restoration initiatives in European open and semi-open lowlands, especially in wetland areas of high conservation value (Gordon et al., 1990; Wallis DeVries, 1998; Carver et al., 2021). Much evidence suggests that these large herbivores are able to substitute their ancestors – wild horses and aurochs – by creating and maintaining spatial heterogeneity (Trepel et al., 2024) and thus contributing to species richness and diversity (Olff and Ritchie, 1998; Stein et al., 2014; Lorimer et al., 2015; Pereira and Navarro, 2015; Gordon et al., 2021; Svenning et al., 2024). Large herbivores shape ecosystems through their movement-, grazing- and resting-behaviours (Liu et al., 2015) and by their habitat use (Martin et al., 2009). This habitat choice is influenced by resource availability (Aarts et al., 2013) and season (Zweifel‐Schielly et al., 2009; Zielke et al., 2019), but it likely primarily depends on the herbivore species and their interspecific interaction, as bovids and equids differ in nutritional and behavioural needs (Olff and Ritchie, 1998; Esmaeili et al., 2021) but show a strong potential for competition due to shared habitat (Menard et al., 2002).

Only few studies of habitat selection of co-existing cattle and horses have been conducted to date (Menard et al., 2002; Lamoot et al., 2005; Thomassen et al., 2023), and these seem to reveal conflicting patterns. For example, Cromsigt et al. (2018) and (Lamoot et al., 2005) showed that horses almost exclusively feed on grasses and thus select grassland habitats, while cattle include a solid amount of woody vegetation in their diet and therefore select wood- and shrubland more often than horses. Thomassen et al. (2023) found high diet richness with low seasonal variability in horses, and less diverse diet yet with increasing summer-time variability in cattle. While these studies reflect considerable differences in habitat use of the two species, others, e.g., Nolte et al. (2014) and Menard et al. (2002), suggest strong competition for grasses between the two species especially in winter, meaning an overlapping habitat choice.

In the first part of our study, we therefore assessed the seasonal variation of space use intensity of semi-wild cattle and horses in a French nature reserve by following the spatiotemporal distribution of the grazers, and contrasted this space use pattern with the habitats on the study site to describe the habitat use of the grazers. Our first study question was: how do the habitat use patterns differ between horses and cattle, and how does such a difference vary between summer and winter?

Not only the habitat use patterns of co-existing cattle and horses are still unclear, but also the effect of this habitat use on plants is not well understood. In a comprehensive review, Bakker et al. (2006) found the effects of large herbivores on plant diversity conflicting, with effects ranging from positive (Belsky, 1992; Collins et al., 1998) through neutral (Stohlgren et al., 1999; Adler et al., 2005) to even negative (Milchunas et al., 1998). This may be because grazers can have either compensatory or additive effects on vegetation through selective feeding on different competing plants, or through consumption of the same plants by multiple herbivore species (Ritchie and Olff, 1999). For example, Schmitz and Isselstein (2020) showed that cattle defoliate grasses more evenly, creating a homogenous sward of grazing-tolerant plants, while horses graze heterogeneously, avoiding foraging in latrine areas and strongly defoliating in other areas. As an outcome, responses of the plant community will result in changes in vegetation structure and diversity (Olff and Ritchie, 1998; Liu et al., 2015).

The response of plant community structure to grazers is believed to be principally determined by functional traits of plants (Díaz et al., 2007). Investigating the response of such functional traits are a “common currency” (McGill et al., 2006) to compare species independently of species identity and across taxa (Reese et al., 2016). Changes in plant functional traits as a function of grazer presence therefore offer generalizable predictions about the effect of rewilding with large herbivores as a means for ecosystem restoration (Atkinson et al., 2024). Yet, as Adler et al. (2005) points out, the challenge is identifying key functional traits and then understanding how differences in their means and variances translate into differences in response to grazing.

Such key traits may include tolerance for nutrient-rich soil, preference for exposure to light, and tolerance of disturbance as predictors of changes in plant community as a function of grazer presence (Milchunas et al., 1988; Augustine and McNaughton, 1998). Further, simple vegetative traits such as plant height and patch cover may describe a general response of herbaceous plant community to large herbivores (Díaz et al., 2001).

Results about how these traits predict plant responses are however often contradicting. Herbivores seem to show opposing effect on plant species with different nutrient-tolerance levels, as they alter rates of nutrient cycling and redistribute nutrient availability (Trlica and Rittenhouse, 1993; Bardgett, 2005). For example, frequently grazed patches may experience “nutrient export”, resulting in a shift towards plant species with lower nutrient demands (Nolte et al., 2014; Schmitz and Isselstein, 2020); but urine and dung input by herbivores may also increase organic matter and nitrogen levels of the soil on intensively visited patches (Harrison and Bardgett, 2008; Peco et al., 2017), leading to an opposite effect: a shift towards nutrient-tolerant species.

Light-preference may similarly result in vegetation composition differences as a function of grazers’ space use, since heterogeneous grazing patterns can lead to spatial heterogeneity of light availability (Bakker et al., 2003). Intense grazer presence at certain areas may reduce competition for light by creating gaps and opening colonization windows in the vegetation, and thus may favour light-demanding plants (Collins et al., 1998). Other areas without frequent grazer visits may however increase competition for light, hosting mainly the few most competitive light-preferent species and an array of shade-tolerant plants as an understory (reviewed in: Marion et al., 2010). Yet, due to the heterogeneous nature of grazing, such patterns are highly variable, and general predictions are hard to draw.

Tolerance to disturbance, such as grazing and trampling (often termed as herbivory) is regarded to be a basic predictor for changes in plant community composition (Augustine and McNaughton, 1998; Díaz and Cabido, 2001; Evju et al., 2009). Herbivory affects plant physiology, morphology, and genetics, thus plants have evolved to avoid or tolerate such disturbances (Trlica and Rittenhouse, 1993). However, the several forms of herbivory impacts and the variation in grazing pressure make predictions about the effects of grazing on different plant species difficult (Reese et al., 2016). Furthermore, the tolerance of plants to grazing depends also on how they interact with neighbouring plants (Hendrickson and Olson, 2006).

As Herrero‐Jáuregui and Oesterheld (2018) emphasised, most of our knowledge of the effect of grazing animals on vegetation structure is based on grazed–non-grazed contrasts, but the effects of a scenario where grazing intensity varies from low to high due to the movement patterns of free-roaming animals are largely unknown. Also, comparatively little attention has been paid to investigating the joint effect of co-existing free-roaming cattle and horses (Liu et al., 2015), especially in nature conservation initiatives.

In the second part of our study, we thus carried out a large-scale macrohabitat survey, and additionally we recorded vegetation characteristics on a microhabitat scale to assess the changes that large herbivores induce in the plant community composition and the physical vegetation structure.

Following Díaz et al. (2007), we assessed the structural responses of the vegetation to answer the question: how does plant height, patch cover and species richness change as a function of the spatiotemporal distribution of the grazers? We were also interested in how plant functional traits of light preference, grazing tolerance and soil-nutrient tolerance changed over four years, including one year before and three years after the introduction of the cattle and horses on our study site.

The distinctive feature of our study lies in the fact that we were able to follow the ecological processes from a “zero state” on an area of a recent ecological restoration initiative. The study site used to be an agricultural area, where, after eliminating the remains of pesticides, the colonisation by vegetation started from bare soil, and where wild-living, free-roaming rewilded cattle and horses were then introduced to fulfil the functional roles of their wild ancestors in regulating natural succession and in contributing to ecosystem processes (Vera, 2000; Lovász, 2022; Schmitz et al., 2023).

Our overall aim was to contribute to a more comprehensive understanding of the seasonal and interspecific variability of habitat use of large herbivores, and of the vegetation functional trait-dynamics as an outcome of this habitat use in a rewilding-based ecosystem restoration area. Our study thus helps to inform management strategies when applying free-roaming horses and cattle in ecosystem restoration, and especially in rewilding initiatives.

## Methods

### Study area

Our study area is situated on an island of the Rhine river and the Grand Canal of Alsace in France, and is part of the national nature reserve Petite Camargue Alsacienne. A total of 100 hectares of the island has been dedicated to an ecosystem restoration project since 2014. Our study site is a 32-hectare area, part of the restored 100-ha area.

In the course of the restoration initiative, former crop fields in the area were transformed into an alluvial environment. The reconstructed landscape comprises open grasslands with scattered shrubs and tree groups, a few gravel sites without vegetation cover, groundwater ponds, and a small creek allowing part of the Rhine water to flow through the island. Remnants of old alluvial forest patches that surround the area remained untouched during the ecosystem restoration. Natural succession from bare ground with gravel areas started with an initial prompt to vegetation growth by seeding and planting indigenous plant species in 2015 (Lachat, 2012). In addition, free-roaming Konik horses and Highland cattle were introduced following the concept of rewilding, i.e., the passive management of ecological succession to restore natural ecosystem processes and reduce the human control of landscapes (Pereira and Navarro, 2015). The cattle and horses arrived between September 2018 and March 2019 and their expected role was to act as a natural disturbance regime (Mackey and Currie, 2000; Vera, 2000) on the restored alluvial environment by creating spatial heterogeneity and thus enhancing biodiversity (Pereira and Navarro, 2015; Carver et al., 2021). The animals have thus been grazing year-round in low grazing intensity (0.3-0.5 animals per ha); human intervention such as supplementary feeding has been restricted so that animals have a natural impact on their environment, especially on the vegetation.

The national nature reserve Petite Camargue Alsacienne granted us permission to conduct our study on this site.

### Grazer spatial distribution

Upon their arrival to the Rhine island, the horses and the cattle were equipped with GPS-collars (Followit, type Pellego) for management purposes of the nature reserve. The collars recorded the position of the grazers every hour, and we used the data of summers and winters from 2019 to 2021 (see Supplementary Material Table 1). We described the space use of the grazers based on these hourly GPS positions. Parts of those data were used in two other studies (Lovász et al, 2021, and Lovász et al 2024), but for better understanding, we give a detailed explanation of the corresponding methods here as well.

During this three-year period, the number of grazers increased from the initial 5 horses and 5 cattle to 7 horses and 7 cattle.

The GPS fixes may show some imprecision because GPS accuracy can be affected by satellite or receiver errors (Hurn 1993), satellite geometry (Dussault et al. 2001), atmospheric conditions, topography, or overhead canopies (Di Orio et al. 2003; Moen et al. 1996). The GPS collars did not record DOP (Dilution of Precision) data and we therefore could not correct for the inaccuracy of the fixes. However, since in an earlier study using a part of the same GPS-collar data (Lovász et al., 2021), only 3.3% of all grazer positions fell outside the fenced area, and since also earlier studies about GPS-error showed marginal average inaccuracy (e.g., Ganskopp and Johnson, 2007), we assumed that inaccuracy would not strongly influence our results; we thus excluded GPS positions erroneously falling outside of the enclosure in our analysis.

The data were downloaded via satellite processing and through the interface of the GPS collar provider (Followit, Lindesberg, Sweden) remotely, without contact to the grazers. The use of GPS collars on both cattle and horses comply with animal welfare requirements and reportedly does not cause disturbance to the animals (Ungar et al., 2005; Collins et al., 2014).

### Macrohabitat data

We carried out visual landcover surveys on the 32-ha study site in the median year of our sampling period (in 2020). We divided the study area into 50×50-meter grid cells, and with the support of a printed map with the projected grid cells, the observer (L.L.) surveyed each grid cell by standing in the midpoint of the cell and visually estimating the percentages of landcover types in the respective grid cell. The data from these surveys were also used in (Lovász et al., 2024).

We distinguished the following landcover categories: trees (trees of ca. ≥ 3 years old), saplings (all growth stages of young trees of ca. < 3 years old), shrubs, meadow, bare ground, and surface of water in the respective grid cell. These macrohabitat-scale habitat characteristics remained relatively stable throughout the 3-year study period.

### Microhabitat data

We sampled herbaceous vegetation yearly between 2018 and 2021 on 80 permanent quadratic plots of 1 m^2^ each. The locations of the plots were defined by the mid-points of the 50-meter UTM-grid cells that we used also for the macrohabitat data. The coordinates of the sample points were determined by using QGIS (version 3.4.4-Madeira). Two months before the first data collection, we marked the sampling plots by digging coloured flat concrete tiles into the ground, to facilitate finding the plots visually in high vegetation and to avoid that the grazers stumble on them. To avoid potential effects of the marking tiles on the vegetation, the closest corner of a sampling plot was placed at a distance of 50 cm from the north-western corner of a tile.

Vegetation sampling (performed by L.L.) took place during the peak growing season, between mid-May and mid-July. For the measurements, we used a modified Daubenmire method (Daubenmire, 1959), by laying a 1×1-meter sized frame onto a sampling plot. Within each plot, the following attributes were measured: plant height (cm); total vegetation cover (%); species (name) of the three most abundant plant species. The variable ‘plant height’ was the average of 20 measurements per plot that were taken along the two diagonals of the 1-m^2^ frame every 10 cm. We measured the total vegetation cover in the plot by visually estimating the percentage of the plot covered by plants (on a continuous scale). The three most abundant plant species in each plot were identified according to visual estimation of their abundance; when identification to the species level was not possible due to missing identification criteria (e.g., in graminoids, the lack of flowers due to grazing), we identified the plant to the genus level.

Using the dataset of Flora Indicativa (Landolt et al., 2010), the following functional-trait attributes were assigned to each species identified in our study area: light preference, soil-nutrient tolerance and mowing/grazing tolerance (see Table 1 for the scales used to describe the plant species).

We estimated how the means of these functional traits of the most abundant species in the 80 sampling plots changed as a function of horse and cattle location density.

**Table 1.** Plant functional trait attributes according to Landolt et al. (2010).

| Scale | Light (L) | Nutrients in the soil (N) | Mowing/grazing tolerance (MT) |
| --- | --- | --- | --- |
| 1 | full shade | very infertile | bad |
| 2 | shade | infertile | little |
| 3 | semi-shade | medium fertile | moderate |
| 4 | light | fertile | good |
| 5 | full light | over-rich | very good |

Because vegetation sampling took place during the summers of the study period, we assessed the effect of average grazer location density on the functional and structural traits of plants in summer only. For each trait, we assessed the response to both horse and cattle location density, and we also estimated average yearly changes.

In addition to the mean light preference of the three most abundant plant species in a plot, we also assessed variation between plots by calculating the standard deviation of light preference among the species.

We also calculated the overall species richness, species (alpha) diversity and relative evenness of the plant community for each year of the study, using the species list arising from taking the three most abundant species identified from all sampling plots. Species richness refers to the sum of all species recorded on the study site in a given year. For calculating species diversity, we used the Shannon-index H’ = - ∑*p*_*i*_ ln(*p*_*i*_), where *p* is the proportion of individuals of the *i* species divided by the total number of species. We computed the relative evenness using the Pielou’s index J = H’/ln(N), where H’ is the Shannon-index and N is the total number of species in our study site. Both indices were measured taking the number of occupied cells per species as if they were the numbers of individuals.

### Statistical analysis

For the macrohabitat analysis, we used the 50×50-m grid cells of the UTM grid over the study area. We calculated the average number of locations in a grid cell across days and individuals within each of the six seasons, i.e., summer and winter in the years 2019 to 2021. We averaged across days and individuals because seasons differed in duration and a different number of animals were tracked in the different seasons (Suppl. Mat. Table 1). The average number of locations in a grid cell was calculated separately for horses, cattle and the sum of the two, i.e. “grazers”. We increased the three variables by half of their minimal non-zero value to get rid of the zero-values before we applied the logarithm transformation.

We analysed the logarithm transformed average number of locations per grid cell using a liner mixed model with season (summer vs. winter), landcover variables and the two-way interactions between season and each of the landcover variable as predictors. The grid cell ID was used as a random factor to account for repeated measures of the same grid cells over the six seasons. We included only five of the six landcover variables because the six sum up to 100% which would cause redundancy in the model if all were included. We decided to drop meadow because that was the most abundant landcover and leaving this variable out produces a set of predictors with the lowest possible correlation. Of all landcover variables, both linear and quadratic terms were included.

We checked the model assumptions (normal distribution, linear relationship, homogeneity of variance) by standard diagnostic residual plots. We further checked for spatial correlation by plotting the residuals in space (bubble plot) and by the semi-variogram.

To analyse how grazer location density influences vegetation structure, we summed grazer locations within each grid cell from 1 January to 1 June of each of four years. During the first year 2018 grazer density was zero. Because not all animals were tracked and the number of tracked animals varied over the five-month period, we corrected that number by dividing it by the number of “animal-days” of tracked animals and multiplied it by the number of “animal-days” of the total number of animals. The total number of horses varied between 5 and 9 individuals (of which 3 to 5 were tracked) and the total number of cattle varied between 3 and 8 individuals (of which 2 to 5 were tracked). In that way, we got for every 50×50m grid cell an expected grazer density that was obtained from the space use of the grazers and the total grazer abundance. We square-root transformed the horse and cattle grazer density variables before we used them as predictors in the analyses of vegetation structure.

We used a normal linear mixed model for analysing each of the six vegetation trait variables: average light preference value, standard deviation of light preference value, average nutrient-tolerance value, average mowing/grazing tolerance value, total cover, average height. We used horse and cattle location density and year as numeric predictors, and the cell ID was used as a random factor to account for the repeated measure of the same cells over the four years. Standard diagnostic residual plots were used to assess normal distribution of the residuals, linearity assumption and homogeneity of variance. We further used semi-variograms to check for spatial correlation and auto-correlation plots to check for temporal correlation.

Data preparation and model fitting was done in R 4.2.2 (RCoreTeam, 2022). For fitting the normal linear mixed models, we used the function lmer of the package lme4 (Bates et al., 2015). We used flat prior distributions for all model parameters to obtain posterior distributions from which we report the 2.5% and 97.5% quantiles as lower and upper limits of compatibility intervals of parameters and fitted values. The function sim from the package arm (Gelman and Hill, 2007) was used to obtain the posterior distribution.

## Results

### Landcover characterization

From the six landcover types (see Fig. 1), meadow cover was the most abundant (42.3%). The relatively high percentage of tree cover (32.6%) was mostly due to an old oak forest in the south-eastern corner of the grazed area, while at other parts of the study site, trees were found only in small patches. The percentage of sapling cover (11.9%) indicated a fast succession of mainly poplar trees. Open bare ground and shrub cover were present in small proportions (4.7% and 2.4%). Water bodies represented 6.2% of all cover types and were all groundwater ponds with stable water levels throughout the year.

**Figure 1.**
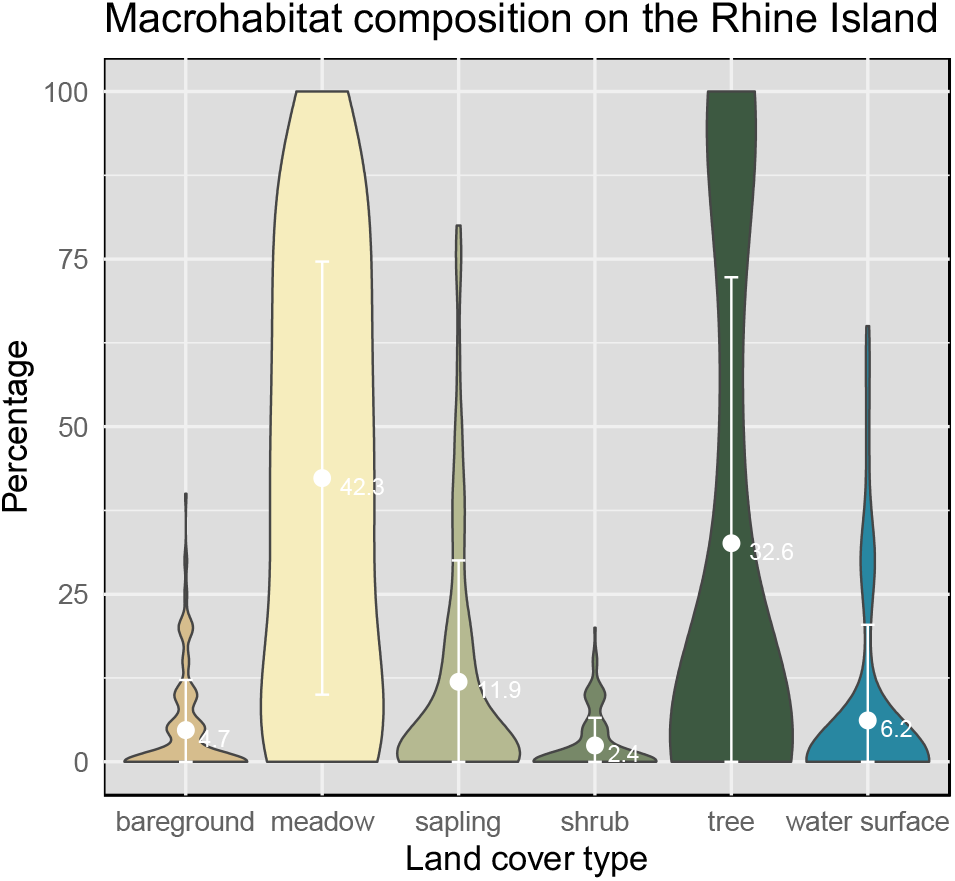
Macrohabitat composition on the Rhine Island. Violin plots show the percentages for each land cover type, with mean percentage (white dot, and value in %) and standard deviation (white error bars) over all grid cells sampled

### Macrohabitat use of horses and cattle

In our study site, Highland cattle and Konik horses used the different habitats in a similar manner, but we found differences in seasonal patterns and the magnitude of the patterns (Fig. 2 and 3, and Suppl. Mat. Fig. 1).

**Figure 2.**
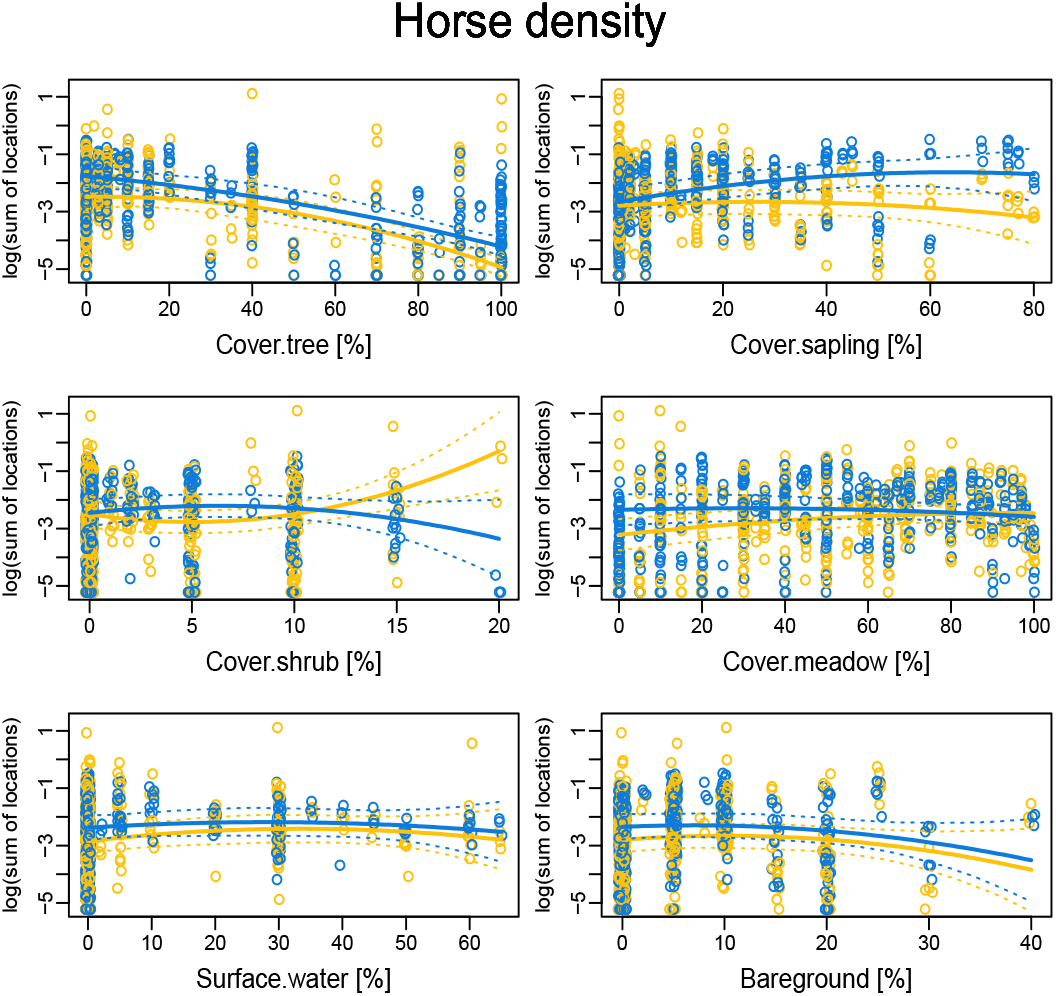
Logarithm of horse location density in relation to the cover variables in summer (orange) and winter (blue). Dotted lines are 95% compatibility intervals

**Figure 3.**
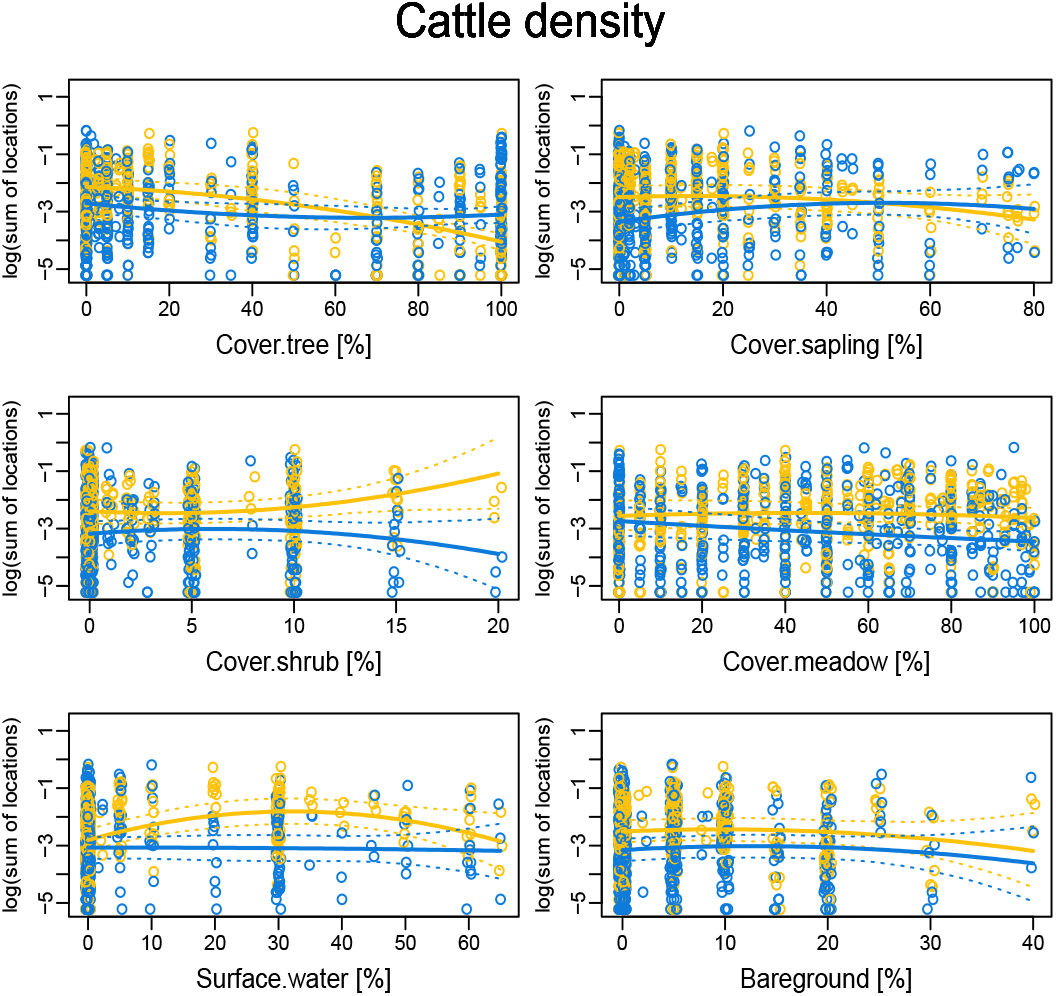
Logarithm of cattle location density in relation to the cover variables in summer (orange) and winter (blue). Dotted lines are 95% compatibility intervals

Both grazer species were similarly less frequently present in areas with high than with low tree cover in summer, but in winter cattle densities outnumbered horse densities in forested grid cells (Fig. 2 and 3).

Areas with higher sapling cover were increasingly used in winter by horses but not so much by cattle. In summer, sapling-covered areas seemed not to be very important habitats for either of the grazer species, as regression lines show a likely negative relationship (Fig. 2 and 3, but note that the compatibility intervals (Amrhein and Greenland, 2022) would allow a slight positive relationship too).

Patterns were clearer regarding the use of grid cells with shrubs both in case of cattle and horses (Fig. 2 and 3), with a preference for such shrub-covered habitats in summer. However, note that shrub cover was rather patchy in our study site, with a maximum of 20% cover by grid cell, and an average of 4.7% of the total area (Fig. 1), therefore locations of animals in these habitats may be a consequence of selecting neighbouring habitats (which were often meadows, since meadow was the most abundant cover variable in our dataset).

Meadows – i.e., habitats of extensive grass cover – were used rather evenly by both grazer species, except in winter, when cattle densities decreased as meadow-cover increased. This refers to a preference towards habitats other than only meadow-covered areas in winter. In summer, the location densities of cattle were quite similar in all scales of grass-cover, possibly meaning a preference for meadows compared to other cover variables, yet without a strong trend. Horses on the other hand were aggregating on grid cells with the highest meadow-cover percentages in summer.

Grid cells with average percentages of waterbodies were used by cattle more often in summer than in winter, while we observed horses in similar densities in both seasons in areas with water presence.

Unsurprisingly, neither grazer species selected areas with extensive bare-ground percentages, both cattle and horses showed a clear negative relationship to bare-ground.

### Grazer spatiotemporal densities and plant communities on a microhabitat scale

In the 80 sampling plots, the mean number of light-preferring plant species (Light mean; Fig 4, Table 2) did not seem to change at areas with different intensity of occurrence (i.e., GPS-location density) of horse and cattle (effect size: 0.00, CrI: 0.00 – 0.01 for both horses and cattle). At the same time, throughout the 4 years of the study, there seemed to be a slight decrease in species with higher light preference, but the effect was not strong (effect size: -0.03, CrI: -0.07 – 0.01).

**Table 2.** Response of plant functional and structural traits to grazing by horses and cattle. Effect sizes and compatibility intervals.

|  | Interc. | Int.lwr | Int.upr | horse | horse.lwr | horse.upr | cattle | cattle.lwr | cattle.upr | year.z | year.lwr | year.upr |
| --- | --- | --- | --- | --- | --- | --- | --- | --- | --- | --- | --- | --- |
| Light.mean | 3.38 | 3.30 | 3.46 | 0.00 | 0.00 | 0.01 | 0.00 | 0.00 | 0.01 | -0.03 | -0.07 | 0.01 |
| Light.SD | 0.40 | 0.33 | 0.47 | 0.00 | -0.01 | 0.01 | 0.01 | 0.00 | 0.02 | -0.02 | -0.06 | 0.02 |
| N.mean | 3.02 | 2.90 | 3.15 | 0.00 | -0.01 | 0.01 | 0.01 | -0.01 | 0.02 | -0.07 | -0.13 | 0.00 |
| MV.mean | 2.88 | 2.73 | 3.03 | 0.01 | 0.00 | 0.03 | -0.02 | -0.04 | 0.00 | 0.23 | 0.15 | 0.31 |
| log(height.m) | 2.45 | 2.29 | 2.61 | -0.01 | -0.02 | 0.01 | -0.01 | -0.03 | 0.01 | -0.25 | -0.34 | -0.17 |
| total cover | 70.66 | 64.76 | 76.70 | 0.67 | 0.15 | 1.20 | -0.29 | -0.82 | 0.25 | -0.10 | -2.62 | 2.37 |

**Figure 4.**
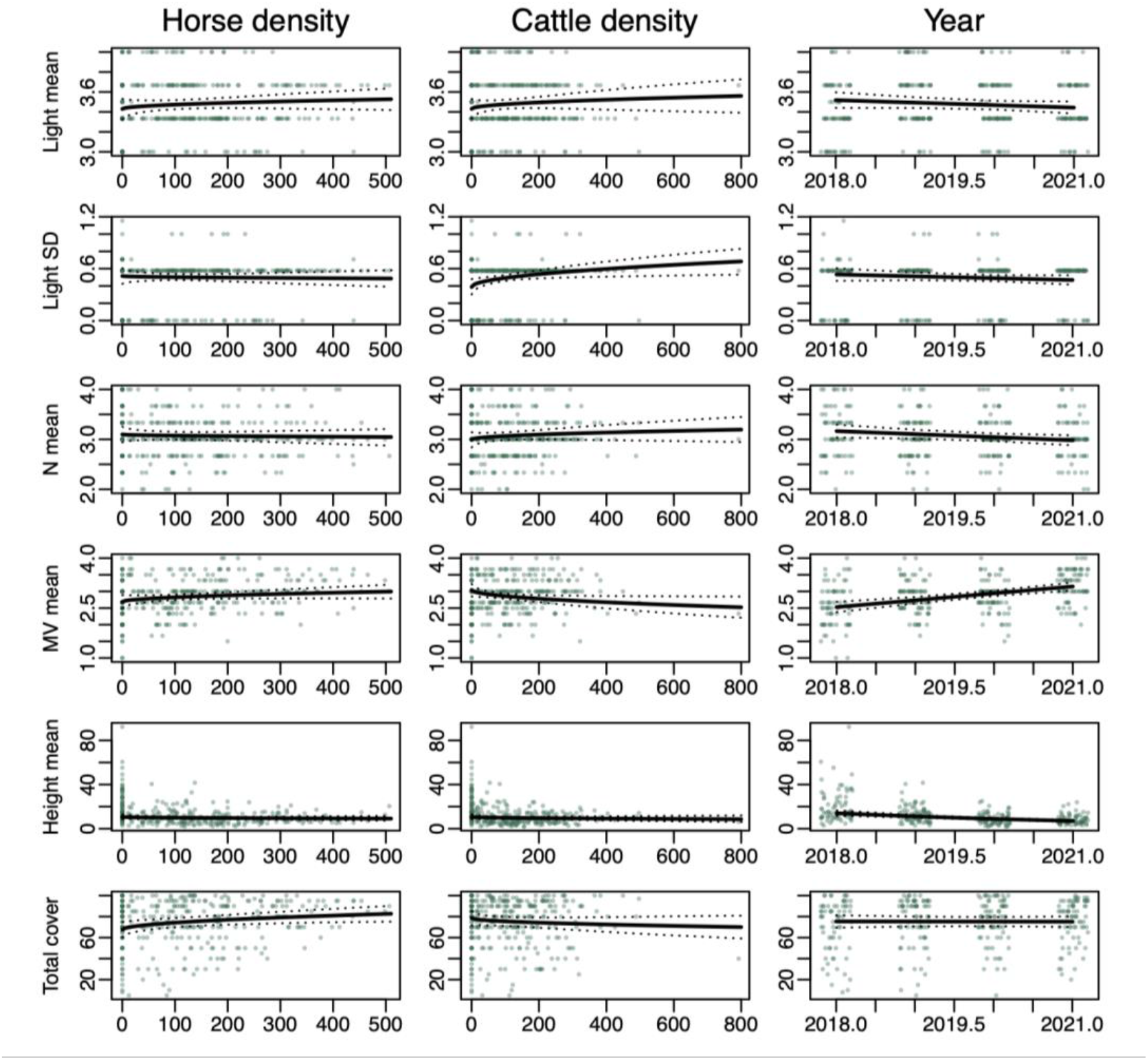
Relationship between horse and cattle location densities and plant functional (mean of light-preference values [Light mean]; standard deviation of light preference values [Light SD]; mean nutrient-tolerance [N mean]; mean of mowing/grazing tolerance values [MV mean]) and structural traits (mean height [cm] and total cover [%]), and the changes of the plant community along these traits throughout the 4 years of the study.

The variation between species of different light preference values (Light SD; Fig 4, Table 2) showed a relatively flat relationship to horse density, but at higher cattle densities, the within-plot variation in light preference among plant species was higher. The long-term general trend was a rather slight decrease in the variation of species according to light preference over the 4 years (effect size: -0.02, CrI: -0.06 – 0.02).

The mean number of plants with nutrient-rich soil preference (N mean; Fig 4, Table 2) did not seem to differ in areas with different intensity of horse presence (effect size: 0.00, CrI: -0.01 – 0.01), and their numbers were only slightly higher at higher cattle densities (effect size: 0.01, CrI: -0.01 – 0.02). We found a slight yearly decrease in nutrient-preferring plants (effect size: -0.07, CrI: -0.13 – 0.00).

Grazing-tolerant plant species (MV mean; Fig 4, Table 2) increased with increasing horse densities but decreased with cattle densities. In the course of the four study years, there was a clear increase in grazing-tolerant plant species (effect size: 0.23, CrI: 0.15–0.31).

Mean vegetation height (Fig 4, Table 2) was similar at low and high densities of both horses and cattle. Yet, throughout the years, the mean vegetation height clearly decreased (effect size (log): -0.25, CrI: -0,34 – 0.17).

The average of total vegetation cover seemed to increase in areas where horse density was higher but the direction of the effect in case of cattle location densities indicated a potentially negative relationship (although compatibility intervals might also allow for the opposite direction of the effects, Fig. 4.). However, throughout the four years of the study period – including the first year without grazer presence – the average total cover seemed to stay rather stable (effect size: -0.1, CrI: -2,62 – 2.37). This may suggest that the effects of horses and cattle potentially balance each other out on the overall vegetation cover.

The total number of species decreased after grazers were introduced (2019), and this trend continued until the third year of grazing (2021), when species richness slightly increased again. Similarly, the Shannon-index showed a small decrease in diversity until 2021, when diversity increased again. The Pielou’s index of evenness however almost did not decrease from 2018 to 2019, and after a small decline, it slightly increased in the last year of the study (Table 3.)

**Table 3.** Diversity of vegetation on the study site in the different years: Species richness (Nr of species), Shannon diversity index and Pielou evenness.

| Year | Nr of species | Shannon index | Pielou evenness |
| --- | --- | --- | --- |
| 2018 | 51 | 3.352321 | 0.8526119 |
| 2019 | 46 | 3.263625 | 0.8524238 |
| 2020 | 42 | 3.099439 | 0.8292438 |
| 2021 | 43 | 3.138654 | 0.8344821 |

## Discussion

### Macrohabitat use of horses and cattle

Our investigation of the habitat use of co-existing free-roaming rewilded horses and cattle revealed overall similar habitat use, but with seasonal and, to some degree, directional differences. In light of our results, we assume that these two species share feeding niches to a certain level and may compete for certain resources, as for instance the findings of Scasta et al. (2016) suggest in western North American rangeland habitats, or as Menard et al. (2002) found in European wetlands. However, since overall grazing pressure remained low in our study site –0.3-0.4 animal per hectare –, resource availability stayed high enough throughout all seasons, therefore strong competition did likely not occur.

During summer, both grazer species exhibited low location densities in areas with high tree cover, but cattle were found in forested areas more frequently than horses. Especially in winter, cattle sought out grid cells in densely forested areas, while horses stayed away from high-percentage tree cover, similarly as for example Menard et al. (2002) and Cromsigt et al. (2018) found. The use of forested habitats by cattle in winter on our study site is in contrast with some earlier findings that showed the lowest probability of woodland selection by cattle in autumn and winter and the highest in spring and summer (Lamoot et al., 2005). On the other hand, the work by Pratt et al. (1986) detected a pattern similar to what we found, and assumed the shelter-seeking behaviour of cattle as a reason for searching woodlands in winter. This may have also played a role in our study area. However, on our study site, the reason why Highland cattle might have searched for forested areas in winter could have been their capacity to reach higher branches of trees with their horns and by bending them down, they could eat the leftover leaves and bark of the branch (L.L. personal observation). Previous studies (Putman et al., 1987; Bokdam et al., 2003) investigated the habitat use of horn-less cattle breeds, which may have been the cause of finding overwhelmingly higher grassland than woodland use by cattle. Our results may thus indicate that conservation grazing involving cattle breeds with long horns, such as Highland cattle, may facilitate the suppression of woody encroachment.

Horses, on the other hand, used areas with higher sapling cover during winter instead of old woodlands. Such habitat preference in free-roaming horses was, to our knowledge, so far not described. An earlier work found increased tree or shrub consumption of horses in winter (Thomassen et al., 2023), but in their study site, sapling-covered areas were not present in similarly high percentage as in our site. It seems therefore that in winter, upon availability, horses rather choose to feed on young trees than on older trees, by eating twigs and debarking thicker stands (L.L. personal observation). Since cattle only showed a relatively neutral relationship to sapling cover in winter, our results may refer to a possible niche separation by the two species.

Water-bodies on the Rhine Island did not seem to have a strong effect on horse location densities, but cattle seemed to search for specific grid cells with water-cover in them. This may be because – according to our personal observations (L.L.) – Highland cattle often cool themselves in warm weather in ponds, and likely also spend much time standing in water to avoid insect harassment. Such behaviour is a natural response to environmental conditions and was not observed in studies about domestic cattle (Senft et al., 1985). Our result may therefore indicate that, to fulfil the natural needs of grazers in nature conservation areas, the presence of large ponds is desirable over drinking wells.

As for the use of meadow habitats, horses seemed to be present constantly throughout winter regardless of the percentage cover of meadows by grid cell, while in summer, horse location density was larger in grid cells with very high (80-100%) meadow cover. Our results are in line with other studies showing that horses prefer grassland areas year-round (e.g., Girard et al., 2013; Scasta et al., 2016).

Cattle, on the other hand, were not found more often in habitats with high percentage of grass cover in summer, even though the availability of meadows was high in our study site (42,3% in total), and even though this species is also considered to be a grazer rather than browser and to select for grasslands more often than woody areas (Pratt et al., 1986). However, as it is theoretically assumed and also recently shown by Thomassen et al. (2023), in co-existing herbivore communities, niche separation between cattle and horses may lead to extended consumption of woody vegetation by cattle. Our results are in line with this theory. Such patterns likely make it possible that the two grazer species may co-exist in natural circumstances, and at the same time have a substantial effect on the vegetation composition (Vermeulen, 2015), facilitating the aims of rewilding initiatives.

### Grazer spatiotemporal densities and plant communities on a microhabitat scale

Our analysis of vegetation functional and structural traits in relation to grazer location density showed different effects of horses and cattle, which seemed to either cancel out or lead to different directional changes when looking at the long-term effects throughout the years.

In the apparent response of the functional traits of plants to grazers, we did not observe strong differences between herbivores and between years, except in the trait of grazing tolerance. Only horse location densities seemed to be related to a higher number of grazing-tolerant plants, while lower numbers of these plants were found at higher cattle densities. This may be because free-roaming cattle are known to spend only about 30% of their daily budget with grazing (Van Rees and Hutson, 1983; Tofastrud et al., 2018), therefore the areas where their location densities accumulated in our study site were likely not at feeding but rather at resting/ruminating areas. Resting and ruminating do not involve defoliating disturbance on the vegetation, which can explain why cattle-presence was negatively correlated with grazing-tolerant plant abundance. In contrast, since horses graze about 50-70% of their daily time budget (Gudmundsson and Dyrmundsson, 1994; Houpt, 2005), high horse densities in our dataset likely indicated feeding activity (rather than only trampling) by horses and thus defoliation disturbance on the vegetation. This may also mean that in our dataset, the classification of plants by Landolt et al. (2010) as grazing/mowing tolerant tended to include tolerance to defoliation rather than tolerance to trampling. In view of this, it is not surprising that the average number of grazing/mowing tolerant plants increased throughout the years (Pakeman, 2004; Evju et al., 2009), also explaining why horse-density had a stronger influence on such plants than cattle-density.

Less clear were the effects of the herbivores on light- and nutrient-preferent plants.

The mean number and also the within-plot variation of light-preferent species seemed to be mainly independent from densities of both horses and cattle.. The relatively flat regression line of the correlations of both the mean and the standard deviation of light preference and grazer densities throughout the four years of the study may indicate a landscape-scale stability over years (Briske et al., 2003).Yet, the slight increase in the within-plot variation of light-preferring plants at higher cattle densities may refer to heterogeneity on the patch-scale and on the short term (Oñatibia et al., 2018; Li et al., 2021).

A similar trend seems to be apparent in the nutrient-preference trait of plants. The nutrient distribution capacity of large herbivores (Doughty et al., 2016) may have resulted in altered nutrient availability in the soil which could lead to differences in nutrient-tolerant species abundance along the gradient of grazer densities (Harrison and Bardgett, 2008), especially in cattle-frequented areas. Yet, the changes across years did not indicate a clear difference and thus may suggest an overall stability of the nutrient preferring plant community.

Structural changes, such as plant height and cover, seemed to be not considerable on the long term. The mean total cover in the 80 sampling plots remained constant over the four years of the study, probably because the percentage of vegetation cover increased with high horse densities, and decreased with high cattle densities. Average height remained relatively constant, with a small mean decline of approximately 10 cm-s between the first and the last year of the study. The mixed-species grazing by horses and cattle therefore seemed to have a conserving effect on plant cover and vegetation height, indicating that such grazing effectively prevents vegetation overgrowth, but at the same time, does not destroy plant physical structure. Our results indicate that, contrary to some earlier findings (e.g., Eldridge et al., 2019) and the concerns of conservation managers, multi-species grazer assemblages in close-to-natural (i.e. 0.3-0.4 animals per ha) densities do not have a destructive impact on herbaceous plants, but are rather an effective tool in conservation management, in line with recent findings (Bonavent et al., 2023).

Our estimation of species richness, diversity and evenness indices suggest slight vegetation compositional changes, likely as a response to grazing. The limitation of our estimation may be that we used only the three most abundant species rather than the entire species pool in each of the 80 sampling plots as a proxy to calculate the diversity indices, the results nevertheless seem to reflect a realistic trend. We detected a small decrease in all indices at the beginning of the grazer introductions, which may be a result of the sensitivity of some of the species (Olff and Ritchie, 1998) that were sown by the management (L. Merckling, 2022, pers. com.) to establish the local species pool on a ‘zero-state’ environment. Further, after introducing the herbivores, a visible decrease of invasive plant species (e.g., *Solidago gigantea*) was observed (L. Merckling, 2022, pers. com., and L.L. personal observation) on the grazed area, which may be due to the effectiveness of grazers in reducing certain alien species to the advantage of local species, similarly as Firn et al. (2013) and Marchetto et al. (2021) found. These two factors therefore might have decreased the total number and diversity of species shortly after the grazer introduction, until it again started slightly increasing in the third year of the grazing management, possibly by species that are more resistant to local conditions, as proposed by Olff and Ritchie (1998). An opposite pattern was shown by Kimball and Schiffman (2003) and McIntyre et al. (2003) but only for cattle grazing and in American and Australian environments, where conditions substantially differ from the European context. Further research could thus investigate the background of such effects in European mixed-grazer assemblages.

## Conclusion

This study fills a gap in previous knowledge by linking the habitat use of rewilded cattle and horses with their effect on functional traits and structure of vegetation, since studies so far mainly investigated one of these subjects but not each of them. The results extend our current understanding of how these animals influence the plant community, by showing complementary habitat use and conserving effects on plant functional and structural traits. We conclude that the varying spatiotemporal distribution of these grazers likely involves changes on the patch scale but vegetation stability on the landscape scale, which is known to facilitate ecosystem functions. Our work may therefore serve as an important source for conservation managers: the results support nature conservation involving mixed species of large herbivores along rewilding principles.

## Acknowledgements and funding

We thank the team of the Réserve Naturelle Petite Camargue Alsacienne for making it possible to conduct our research in the nature reserve. We are grateful for Heiner Lenzin for his support in plant species determination in the initial phase of the study. This work was supported by the Fondation de bienfaisance Jeanne Lovioz, the Foundation Emilia Guggenheim-Schnurr, the Ornithologische Gesellschaft Basel, the Swiss Association Pro Petite Camargue Alsacienne, the Foundation Wolfermann-Nägeli, the Foundation Frey-Clavel, and the MAVA Foundation. The funders had no role in study design, data collection and analysis, decision to publish, or preparation of the manuscript.

## Author Contributions

*Lilla Lovász* conceived, designed and performed the experiments, analyzed the data, prepared figures and/or tables, authored and/or reviewed drafts of the article, and approved the final draft.

*Fränzi Korner-Nievergelt* conceived and designed the experiments, analyzed the data, prepared figures and/or tables, authored and/or reviewed drafts of the article, and approved the final draft. *Valentin Amrhein* conceived and designed the experiments, authored and/or reviewed drafts of the article, and approved the final draft.

