## Supplementary material for "Habitat use of rewilded horses and cattle as related to the functional and structural composition of plant communities in a European restored wetland ecosystem"

Manuscript title:

Table 1. *Number of horses and cattle tracked during the six study seasons*

| year | season | dates | duration (day) | nr of horses tracked | nr of cattle tracked |
|---|---|---|---|---|---|
| **2019** | winter | 9.1. - 28.2. | 51 | 5 | 3 |
| **2019** | summer | 24.6. – 31.8. | 69 | 5 | 2 |
| **2020** | winter | 12.1. - 29.2. | 91 | 5 | 5 |
| **2020** | summer | 1.6. – 31.8. | 92 | 5 | 5 |
| **2021** | winter | 12.1. – 28.2. | 90 | 4 | 2 |
| **2021** | summer | 28.6. – 31.8. | 65 | 3 | 2 |

---

Figure 1. *Logarithm of grazer location density (sum of both cattle and horses) in relation to the cover variables for summer (orange) and winter (blue). 95% compatibility intervals are given as dotted lines*

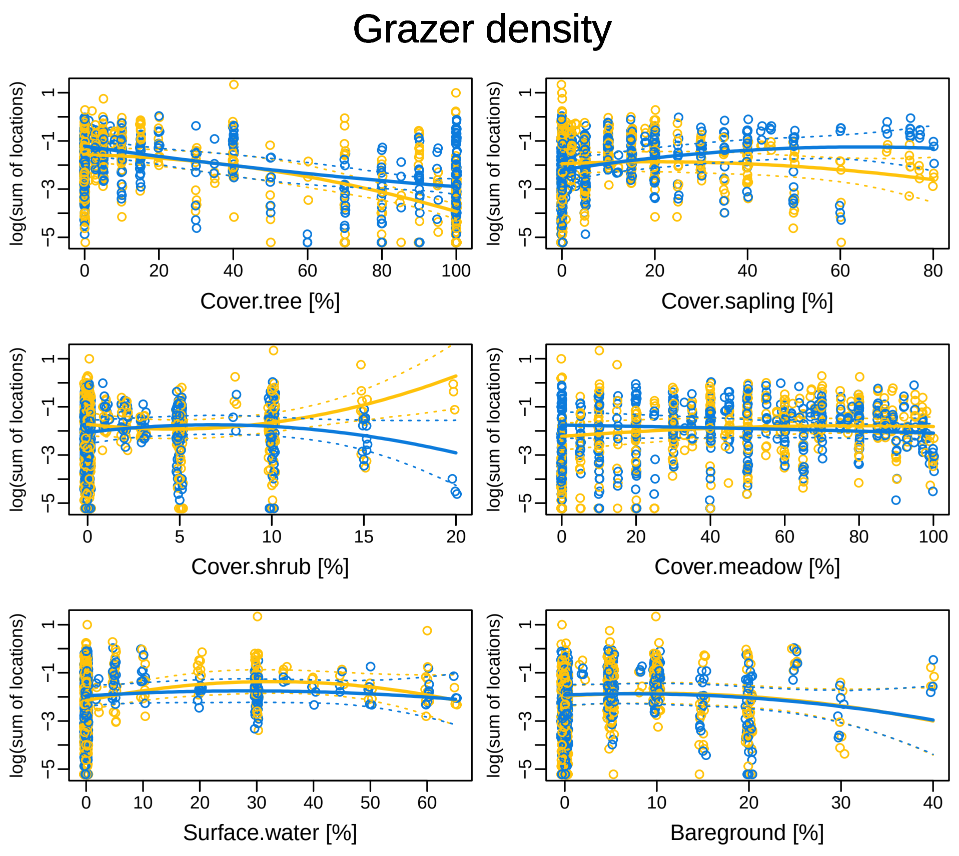
